# Autologous humanized PDX modeling for immuno-oncology recapitulates the human tumor microenvironment

**DOI:** 10.1101/2022.08.19.503502

**Authors:** Michael Chiorazzi, Jan Martinek, Bradley Krasnick, Yunjiang Zheng, Keenan J Robbins, Rihao Qu, Gabriel Kaufmann, Zachary Skidmore, Laura A Henze, Frederic Brösecke, Adam Adonyi, Jun Zhao, Liang Shan, Esen Sefik, Jacqueline Mudd, Ye Bi, S Peter Goedegebuure, Malachi Griffith, Obi Griffith, Abimbola Oyedeji, Sofia Fertuzinhos, Rolando Garcia-Milian, Daniel Boffa, Frank Detterbeck, Andrew Dhanasopon, Justin Blasberg, Benjamin Judson, Scott Gettinger, Katerina Politi, Yuval Kluger, A Karolina Palucka, Ryan Fields, Richard A Flavell

## Abstract

Interactions between immune and tumor cells are critical to determining cancer progression and response. In addition, preclinical prediction of immune-related drug efficacy is limited by inter-species differences between human and mouse, as well as inter-person germline and somatic variation. Here we develop an autologous system that models the TME in individual patients. With patient-derived bone marrow, we engrafted a patient’s hematopoietic system in MISTRG6 mice followed by patient-derived xenograft (PDX) tissue, providing a genetically matched autologous model. We used this system to prospectively study tumor-immune interactions in solid tumor patients. Autologous PDX mice generated innate and adaptive immune populations; these cells populated the TME; and tumors from autologously engrafted mice grew larger than tumors from non-engrafted littermate controls. Single-cell transcriptomics revealed a prominent VEGF-A signature in TME myeloid cells, and inhibition of human VEGF-A abrogated enhanced growth, demonstrating the utility of the autologous PDX system for pre-clinical testing.

## Main Text

The immune milieu within tumors, consisting of diverse cell types including adaptive immune cells as well as macrophages, dendritic cells, natural killer and other innate immune cells, is critical to determining cancer outcome, be it progression or regression.^1^ Macrophages especially can have pro- and anti-growth properties within the tumor microenvironment (TME) in various cancers.^2-4^ However, the immune TME has been challenging to model, owing to inherent inter-species differences.^5-9^ While advances in humanized mice have expanded the repertoire of human immune cells that can repopulate immunodeficient mice, the hematopoietic stem and progenitor cells (HSPCs) used for transplantation have been largely limited to those derived from fetal or neonatal stem cell donors. Thus, these mice do not reflect the systemic immune cell composition of adult humans and remain inadequate models for the majority of human cancers. The ability to pre-clinically model an individual adult cancer patient, capturing the unique features of an individual such as germline genetic determinants of immune function and somatic tumor heterogeneity, is critical to advancing our understanding of interpersonal differences in tumor progression and response to cancer therapies.

MISTRG mice, in which the **M**-CSF (CSF1), CSF2/**I**L3, **S**IRPA, **T**HPO genes were humanized on a **R**AG2^-/-^, IL2R**G**^-/-^ immunodeficient background, generate functional human monocytes, tissue macrophages, alveolar macrophages, and natural killer (NK) cells in a profile more similar to humans than other models, affording an opportunity to model multiple human immune cell types that interact with tumor cells in a human TME.^10-12^ However, efficiency of engraftment in most mouse strains and even the relatively efficient MISTRG mouse precludes the routine engraftment of adult HSPCs. Here, we show that the MISTRG6 strain is a much more adept host recipient of human HSPCs and demonstrate the utility of this system to study the TME in adult solid tumor patients by modeling tumor-immune interactions in a fully genetically matched, autologous manner. We found that humanization of the IL6 locus significantly improved human hematopoietic engraftment of MISTRG6 mice compared with prior models, allowing efficient modeling of individual patients’ tumor-immune interactions using low numbers of HSPCs obtained prospectively from bone marrow (BM) aspirates. Patient-derived xenograft (PDX) tumors grown in mice engrafted with autologous CD34^+^ HSPCs displayed infiltration of patient immune cells of both innate (e.g., macrophage) and adaptive (e.g., T cell) immune subtypes. The transcriptional signatures of the latter imply activation and exhaustion in the TME, whereas the signatures of the former suggest pro-tumor production of VEGF-A in the TME as a major determinant of PDX progression in this model.

## Results

### MISTRG6 displays enhanced proportions of human hematopoietic cells, including innate immune cell types

Given the divergence between human and mouse IL-6 protein (41% identity) and to improve support of human HSPCs, we used the MISTRG6 mouse that bears the human IL6 gene in place of the murine gene in the same chromosomal location, thereby preserving the surrounding regulatory sequences for IL6 control.^13, 14^ When intrahepatically engrafted with equivalent numbers of CD34^+^ cells from human fetal liver (FL), neonatal cord blood (CB), adult mobilized peripheral blood (MPB), or adult bone marrow (BM), MISTRG6 mice harbored greatly increased human hematopoietic cells as a proportion of total hematopoietic cells in peripheral blood compared with NOD-scid-gamma (NSG) and MISTRG mice (Figure 1A). Levels of human hematopoietic cells were also higher in most tissues, including BM, liver, and lung (Fig. 1B), with the greatest differences being between MISTRG6 and NSG hosts. No significant difference was detected in spleen across strains (Fig. 1B). Moreover, we found that MISTRG6 mice could be engrafted with as few as 1,000 human HSPCs, arguably 100x more efficient than other models, and achieve robust hematopoietic transplantation after 10-12 weeks (Fig. 1C), indicating the efficiency of this strain in supporting the growth of hematopoietic cells.

**Figure 1:**
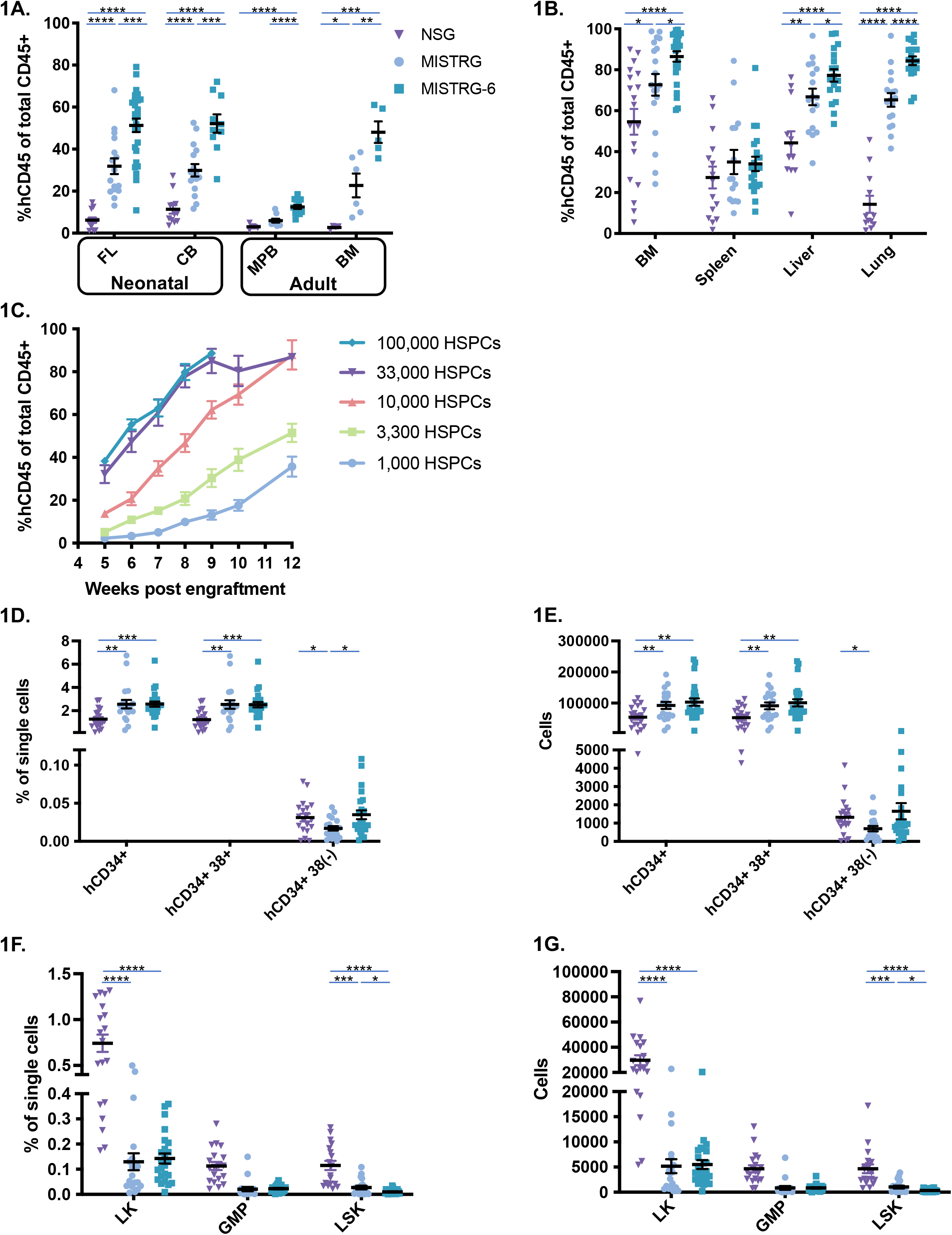
Humanization of the IL-6 locus enhances human hematopoietic engraftment in MISTRG6 mice. A. Human hematopoietic engraftment (percent of hCD45^+^ cells as a proportion of total hCD45^+^ and mCD45^+^ cells) in peripheral blood of NSG (triangles), MISTRG (circles) and MISTRG6 (squares) mice engrafted with equal numbers of CD34^+^ HSPCs from indicated sources; * p<0.05, ** p<0.01, *** p<0.001, **** p<0.0001, unpaired parametric t-test; bars indicate mean and S.E.M.); each dot represents a single mouse. B. Human hematopoietic engraftment (%hCD45^+^ of total CD45^+^) in indicated tissues of NSG, MISTRG and MISTRG6 mice engrafted with equal numbers of CD34^+^ HSPCs. C. Longitudinal analysis of human CD45^+^ cells in peripheral blood of MISTRG6 mice engrafted with varying HSPC numbers as indicated. D. Percentage among single cells of hCD34^+^, hCD34^+^hCD38^+^, and hCD34^+^hCD38^(-)^ cells detected in BM of NSG, MISTRG and MISTRG6 mice engrafted with equal numbers of hCD34^+^ HSPCs. E. Absolute numbers of cells in BM of mice in (D). F. Percentage among single cells of mouse LK, GMP and LSK cells detected by flow cytometry in BM of NSG, MISTRG and MISTRG6 mice engrafted with equal numbers of CD34^+^ HSPCs. G. Absolute numbers of cells from BM of mice in (F).

To better elucidate the mechanism responsible for this enhanced human engraftment, we enumerated human and mouse hematopoietic progenitors in BM of NSG, MISTRG, and MISTRG6 mice. This revealed that human progenitors, including CD34^+^ and CD34^+^CD38^+^ cells, were significantly increased in both frequency and absolute numbers in MISTRG and MISTRG6 mice compared with NSG mice (Fig. 1D-E), and that the mouse hematopoietic lin^(-)^cKit^+^ (LK) and lin^(-)^Sca1^+^cKit^+^ (LSK) progenitor populations were significantly diminished, while granulomonocytic progenitor (GMP) cells were not significantly different (Fig. 1F-G).^15^ In addition, MISTRG6 mice displayed more human hCD34^+^CD38^-^ and fewer mouse LSK cells compared with MISTRG. Collectively, these findings suggest that the enhanced hematopoietic engraftment observed in MISTRG6 is, in part, a consequence of increased human progenitor frequency and reduced mouse progenitor competition.

Humanization of the CSF1 locus in MISTRG enables development of human myeloid lineage cells (CD33^+^) with robust functionality.^10-12^ As expected, this property was preserved in the MISTRG6 model, regardless of human HSPC source or tissue assayed (Supplementary Fig. 1A-F). MISTRG6 mice had a significantly increased proportion of hCD33^+^ myeloid cells than NSG mice in peripheral blood when engrafted with FL- or MPB-derived CD34^+^ cells, with concomitant decrease in frequency of hCD19^+^ B cells (Supp. Fig. 1A, B). In addition, when engrafted with MPB-derived CD34^+^ cells, hNKp46^+^ NK cells were present in equal proportions in MISTRG and MISTRG6 mice but were lacking in NSG mice (Supp. Fig. 1B). These trends persisted in tissues, with CD33^+^ myeloid cells being significantly increased in MISTRG and MISTRG6 mice in spleen, liver, and lung (Supp. Fig. 1C-F). In BM, spleen, liver, and lung, MISTRG6 mice had significantly larger proportions of human NK cells and hCD66b^+^SSC^hi^ granulocytes as well as fewer B cells than NSG mice. T cell proportions did not differ significantly between the strains (Supp. Fig. 1C-F).

In summary, MISTRG6 is highly efficient at supporting development of human hematopoietic cells, including innate immune cells.

### MISTRG6 allows efficient engraftment of patient-derived HSPCs

Having documented that MISTRG6 better supports development of human hematopoietic cells from adult donors and requires transplantation of fewer cells to do so, we sought to apply this prospectively to model individual patients’ TME. For proof of concept, we initially utilized CD34^+^ cells from G-CSF-mobilized peripheral blood samples that were collected from metastatic melanoma patients enrolled in a dendritic cell vaccine trial.^16^ Consistent with the data in Fig. 1 using HSPCs from healthy adults, this showed that engrafting fewer cells yielded higher levels of human engraftment in MISTRG6 compared with MISTRG mice, and that efficient engraftment was feasible with 100,000 to 300,000 CD34^+^ cells (Fig. 2A). Importantly, this enabled engrafting >30 recipient animals. For example, for patient Mel2, engrafting 400,000 CD34^+^ cells in MISTRG hosts yielded a mean of 14.8% human hematopoietic cells in mouse peripheral blood, while engrafting 120,000 CD34^+^ cells in MISTRG6 hosts yielded significantly increased mean human hematopoietic engraftment (34.7%) (Fig. 2A). Similar results were obtained for patients Mel1 and Mel3, for whom fewer cells achieved greater proportions of human hematopoietic cells in MISTRG6 compared with MISTRG, albeit not achieving statistical significance (Fig. 2A). Nevertheless, these results suggest that the enhanced engraftment and growth in MISTRG6 compared to MISTRG is influenced by the individual donor, since Mel3 was superior to Mel1, even though the proportion of HSPCs transferred was greater for Mel1 than Mel3 (800 vs. 220 for Mel1 and 480 vs. 290 for Mel3).

**Figure 2:**
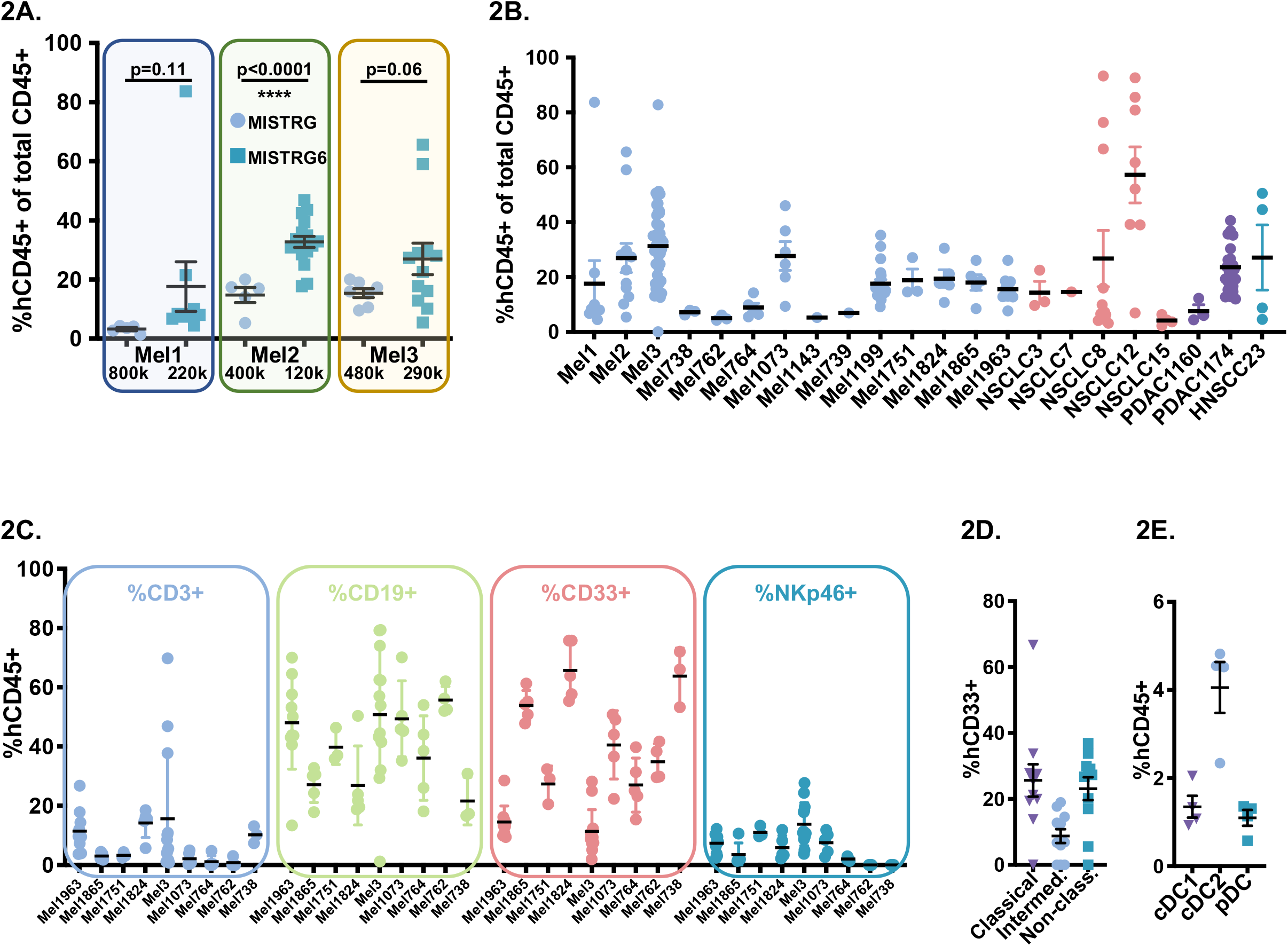
Improved engraftment in MISTRG6 compared with MISTRG mice of HSPCs from solid tumor patients, with human innate and adaptive immune cells represented. A. Analysis of peripheral blood of MISTRG (circles) and MISTRG6 (squares) mice engrafted with HSPCs from the indicated melanoma patients with the HSPC dose shown below; note improved human engraftment levels despite fewer HSPCs introduced in MISTRG6 compared with MISTRG. B. Human hematopoietic engraftment of MISTRG6 mice with HSPCs derived from patients with melanoma (Mel), non-small cell lung cancer (NSCLC), pancreatic adenocarcinoma (PDAC), and squamous cell carcinoma of the head and neck (HNSCC); each dot represents a single mouse. C. Percentage of human T cells (CD3^+^), B cells (CD19^+^), myeloid cells (CD33^+^) and NK cells (NKp46^+^) out of total human hematopoietic cells in peripheral blood of autologously engrafted mice from the indicated patients. D. Proportions of human monocyte subsets (CD14^+^CD16^-^ classical, triangles; CD14^+^CD16^+^ intermediate, circles; CD14^-^CD16+ non-classical, squares) of total CD33^+^ myeloid cells in peripheral blood of Mel1963 autologously engrafted MISTRG6 mice. E. Proportions of human dendritic cell subsets (HLA-DR^+^CD11c^low^CD141^+^ cDC1, triangles; CD11c^+^CD1c^+^ cDC2, circles; CD14^-^CD11c^-^CD303^+^ pDC, squares) of total hCD45^+^ cells in spleens of Mel1963 autologously engrafted MISTRG6 mice.

With successful generation of humanized mice bearing individual patients’ immune systems, we expanded these efforts to prospective collection of BM-derived CD34^+^ cells from patients under active treatment along with tumor tissue from the same patient (i.e., an autologous platform). Under IRB-approved protocols at two cancer centers, we enrolled patients with melanoma (Mel), non-small cell lung cancer (NSCLC), pancreatic adenocarcinoma (PDAC), and head and neck squamous cell carcinoma (HNSCC) to provide BM aspirate, peripheral blood, and tumor tissue at the time of surgery or biopsy. CD34^+^ cells were isolated from BM aspirates using magnetic bead purification and cryopreserved. Viable tumor tissue was utilized to generate PDXs in non-engrafted NSG or MISTRG6 hosts. Overall, 71 patients were enrolled, 46 melanoma, 19 NSCLC, 4 PDAC, 2 HNSCC, ages 22-85, 39% females (Suppl. Table 1). These yielded autologous, immune-reconstituted MISTRG6 hosts from 14 melanoma, 5 NSCLC, 2 PDAC, and 1 HNSCC patients (Fig. 2B).

Autologously engrafted MISTRG6 mice displayed the gamut of human immune cells of adaptive and innate types in peripheral blood at 7 weeks of age (Fig. 2C). Notably, this included CD33^+^ myeloid lineage cells such as CD14^+^CD16^-^ classical, CD14^+^CD16^+^ intermediate, and CD14^-^CD16^+^ non-classical monocytes in peripheral blood (Fig. 2D). Moreover, dendritic cells (DCs), key innate immune cells for initiation of anti-tumor and other immune responses that derive from CD33^+^ cells,^17^ were readily detected by flow cytometry in spleens of autologously-engrafted mice, including human cDC1, cDC2, and pDC populations (Fig. 2E).

### MISTRG6 mice bearing a patient’s hematopoietic cells support autologous PDX growth

Having achieved successful engraftment of patient hematopoietic systems in MISTRG6 hosts, we next subcutaneously introduced the patient’s matched PDX tumor tissue to generate autologously engrafted PDX mice. We monitored tumor growth by caliper measurements in autologously engrafted and non-engrafted (i.e., mice lacking human hematopoietic cells) MISTRG6 littermates, finding that autologous PDX experiments were feasible in MISTRG6 with tumors reaching up to 1000 mm^3^ (Fig. 3A and Suppl. Fig. 2). As expected, PDXs from different patients displayed distinct tumor growth dynamics, with some PDXs growing more or less rapidly. For most patients, tumors grown in autologous HSPC-engrafted hosts were significantly larger than in non-engrafted hosts (Fig. 3A and Suppl. Fig. 2). Harvested xenografted tumors were histologically similar to the parental version and showed hCD45+ hematopoietic infiltration (Fig. 3B). Multicolor immunofluorescence staining of PDX tumors demonstrated that human immune cells, including CD3+ T cells, CD14+ and HLA-DR+ myeloid cells, penetrated deeply into the tumor and co-localized with tumor cells as well as with other engrafted immune cells (Fig. 3C). Indeed, HLA-DR^+^CD14^+^ macrophages and HLA-DR^+^CD14^(-)^ dendritic cells were present, and direct physical interaction between T cells and macrophages was evident (Fig. 3C, bottom panel).

**Figure 3:**
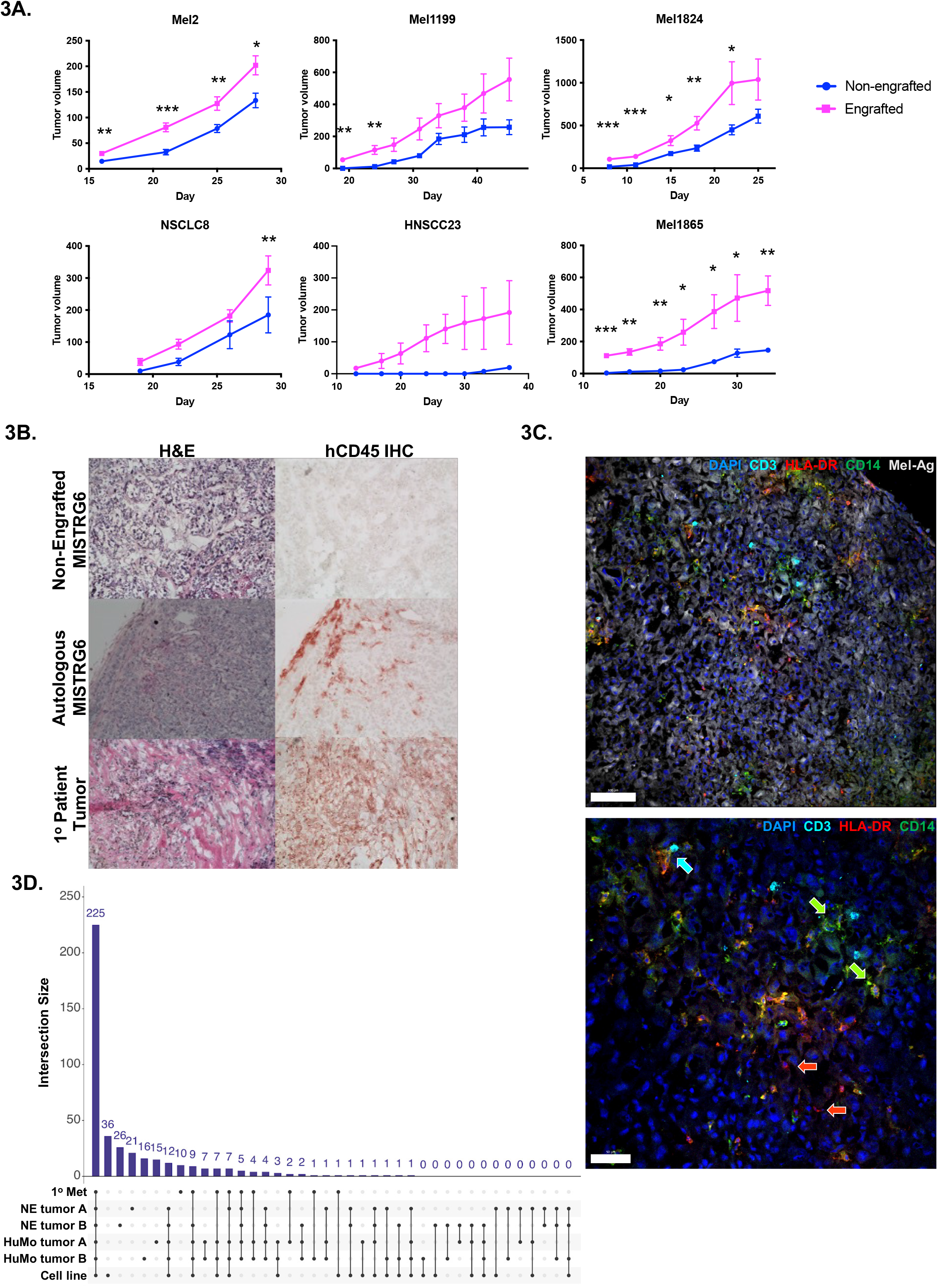
Autologous engraftment enhances PDX growth and human immune cell infiltrate demonstrates tumor-immune interactions. A. PDX growth in MISTRG6 littermates engrafted (magenta) or non-engrafted (blue) with autologous HSPCs from indicated patients (see Suppl. Fig. 2 for comprehensive data). B. Hematoxylin and eosin (left) and hCD45 immunohistochemical (right) staining of tumors harvested from Mel1073 PDX grown in non-engrafted hosts (top), autologously engrafted MISTRG6 hosts and the patient’s primary tumor (bottom). C. Sections of Mel1199 PDX tumor grown in autologously engrafted host stained for human infiltrating immune cells; top panel blue = DAPI, cyan = hCD3, green = hCD14, red = hHLA-DR, white = melanoma antigen, scale bar = 100 um. Bottom panel shows same markers without melanoma antigen staining. Green arrows highlight HLA-DR^+^CD14^+^ macrophages while red arrows highlight HLA-DR^+^CD14^(-)^ dendritic cells. Cyan arrow highlights T cell-macrophage interaction. Scale bar = 50 um. D. Number of somatic mutations (compared with germline reference) shared between the indicated samples: Mel738 patient’s surgical resection sample = 1° Met; PDX tumors from non-engrafted mice (lacking human immune cells) = NE tumor A and B; PDX tumors from mice with autologous engraftment = HuMo A and B; cell line derived from the patient’s tumor = Cell line.

### Whole-exome sequencing reveals preservation of mutations from patient to humanized xenograft

To determine the mutational landscape of tumors generated in MISTRG6 hosts bearing autologous hematopoietic cells and compare to a patient’s tumor as well as those in PDXs grown in non-engrafted mice, we performed whole-exome sequencing (WES) on these samples from Mel738. PBMCs from the patient were used as reference. This analysis indicated that 225 somatic changes were shared between the patient’s surgical resection sample (1° Met), two PDX tumors from non-engrafted mice lacking human immune cells (NE tumor A and B), two PDX tumors from mice with autologous engraftment (HuMo A and B), and a cell line derived from the patient’s tumor (Fig. 3D). 5 additional changes were shared among the tumor samples and absent from the cell line, with 36 additional mutations being specific to the cell line. These data underscore the capacity of the autologous PDX method to recapitulate the somatic heterogeneity that the patient tumor encompasses.

### Autologous MISTRG6 mice display diverse human immune cell populations circulating in the blood and within the tumor, and they recapitulate an immunosuppressive tumor microenvironment

To fully characterize the autologous MISTRG6 model and investigate mechanisms by which autologous human immune cells enhance tumor growth, we performed single cell transcriptomics on hCD45^+^-enriched cells from blood and tumor isolated from Mel1199 and Mel1824 mice (Fig. 4A). After filtering for *bona fide* human cells of high quality, 9,044 cells from blood and 5,559 cells from tumor were analyzed from the two autologous models. All cells were combined, clustered and visualized using UMAP. Examination of cluster-specific gene markers revealed 16 distinct cell subtypes, including 3 myeloid, 2 NK cell, 2 CD8 T cell, 3 CD4 T cell, 2 cycling lymphocyte, 1 B cell, and 3 melanoma cell clusters (Fig. 4B). As expected, these subtypes were differentially represented across tissues (tumor vs blood). For example, monocyte cluster 1 was more highly enriched in blood, while the macrophage cluster was dominant in the TME (Fig. 4C-D).

**Figure 4:**
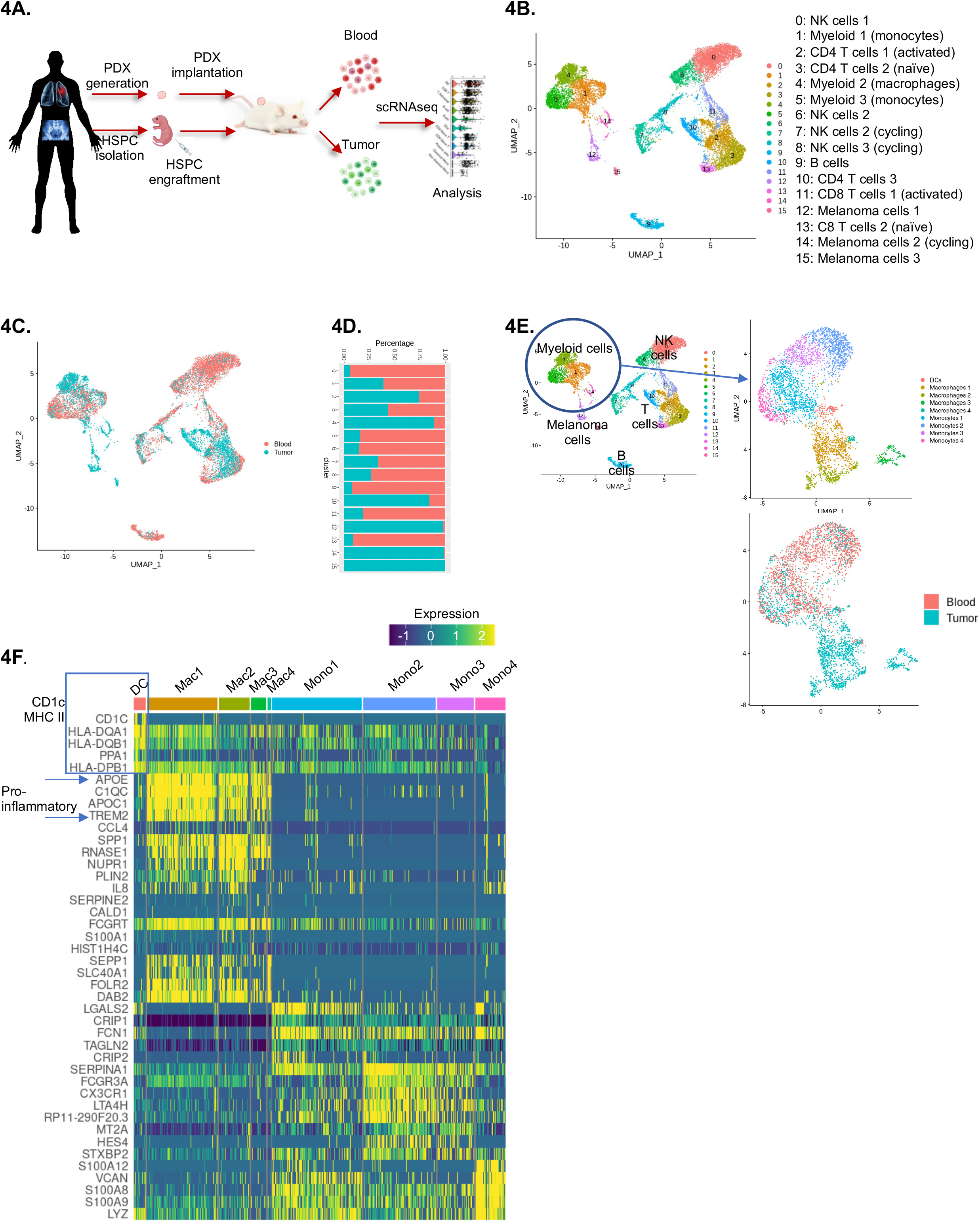
Single cell genomics reveal multiple human immune cell types in tumors and blood of autologous MISTRG6 PDX mice, including innate immune cell types present in the TME. A. Schematic representation of scRNAseq experiments. B. UMAP embedding displaying unsupervised clustering of 14,603 human cells from blood and tumor of autologous MISTRG6 mice. Cell types were identified by marker genes and identities are listed. C. UMAP embedding displaying tissue of origin for cells in A; red cells are derived from blood libraries, blue cells from tumor libraries. D. Proportions of cluster representation in blood vs tumor scRNAseq libraries. E. Re-clustering of myeloid cells reveals sub-structure of 9 clusters including DCs, macrophages and monocytes, in differential tissue representation as indicated in bottom panel. F. Heatmap indicating expression of top differentially expressed genes between each cluster, highlighting presence of human DCs in TME and pro-inflammatory macrophage subtypes.

Given the importance of myeloid cell subtypes in promoting and inhibiting tumor growth, and the unique presence and functionality of these human cells in autologous MISTRG6 models, we reclustered the data to reveal the myeloid subtypes present (Fig. 4E). This revealed 9 distinct clusters including 4 monocyte (distinguished by genes such as LGALS2, SERPINA1, FCGR3A, LYZ), 4 macrophage (defined by APOE, C1QC, APOC1, TREM2), and 1 DC (with high expression of CD1C, FCER1A, FLT3, MHC II genes) cluster (Fig. 4F). As expected, the majority of DC and macrophage cells were detected in the tumor, while most monocytes were present in the blood, reflecting their expected biological homing and plasticity (Fig. 4C-E).

Given the central importance of CD8 T cells as effectors of anti-tumor immune responses, we next characterized the CD8 T cell compartments in autologous models by performing focused sub-clustering of human CD8 T cells (Fig 5A). Comparing CD8 T cells present in blood versus tumor revealed that the most differentially expressed genes (DEGs) found in blood were characteristic of naïve T cells (e.g., LEF1, TCF7, SELL), while genes present in the TME were consistent with activated T cell phenotypes, such as CD69, STAT1, and CXCR4 (Fig. 5B). In addition, sub-clustering revealed 3 distinct CD8 T cell types that included two activated-like populations expressing genes such as CD40LG, XCL1, and XCL2, with one of these populations also expressing an activated/exhausted program typified by expression of PDCD1, LAG3, and GZMA. The third CD8 population expressed naïve-type genes such as LEF1, TCF7, SELL (Fig. 5C). Naïve-like T cells were most highly represented in the blood, while activated and activated/exhausted-like genes were more present in the TME (Fig. 5D).

**Figure 5:**
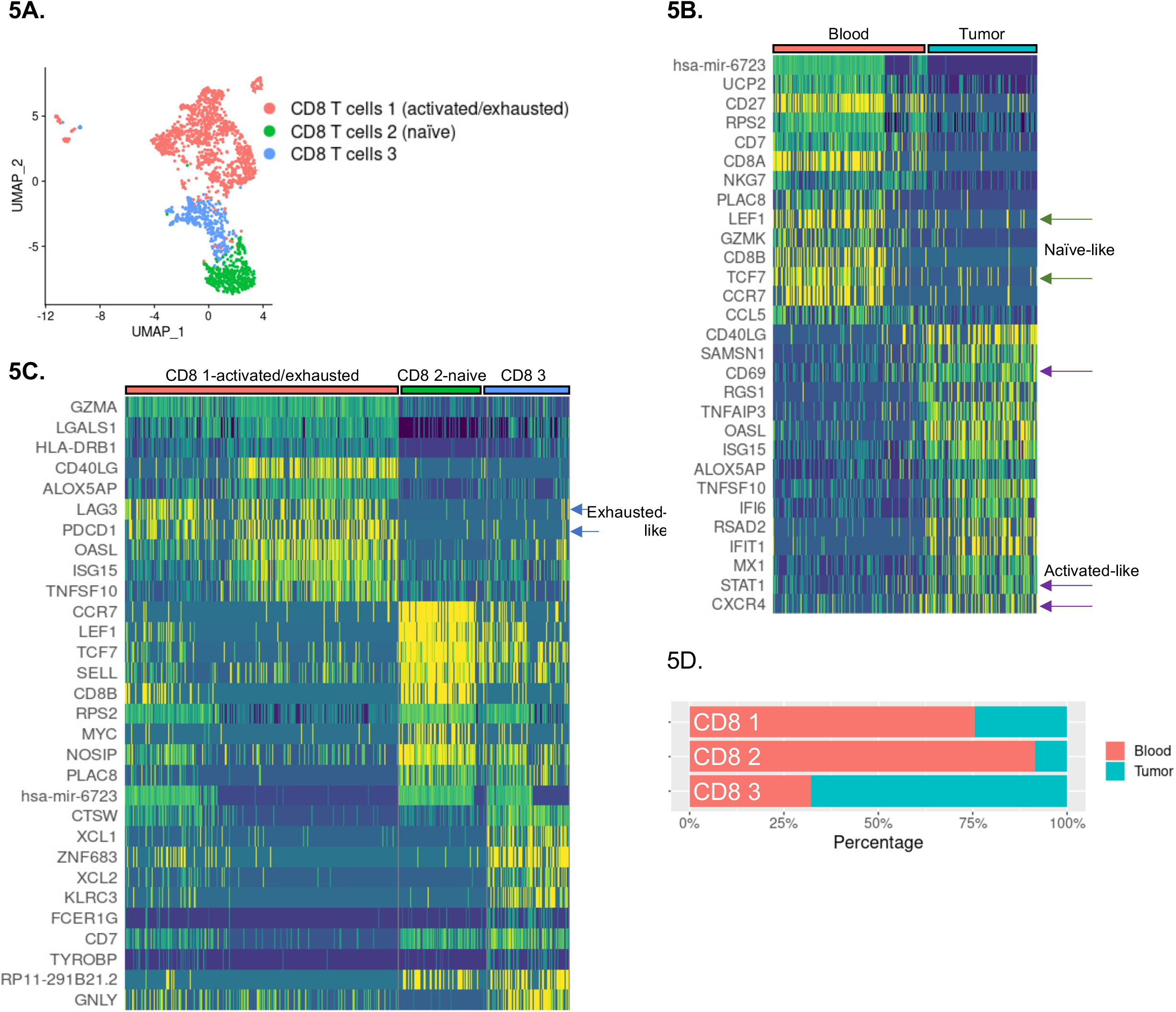
CD8 T cells circulating in the blood of autologous MISTRG6 mice display features of naïve states while those in the TME express markers of activation and exhaustion. A. Re-clustering of CD8 T cells reveals sub-structure of 3 clusters. B. Differentially-expressed genes between CD8 T cells in blood vs tumor display features of naïve (blood) and activated (tumor) states. C. Heatmap indicating expression of top differentially expressed genes between each cluster, highlighting activated/exhausted-like phenotype of CD8 1 cluster. D. Cluster representation of CD8 T cell subclusters in tissues, demonstrating over-representation of activated/exhausted-like CD8 T cells in the tumor microenvironment of autologous mice.

CD4 T cells sub-clustered into 5 distinct groups with divergent localizations between blood and tumor; clusters 1, 3 and 5 were more prevalent in blood, while clusters 2 and 4 were almost exclusively present in tumor (Suppl. Fig. 3A-B). Cluster 1 was characterized by naïve T cell genes, while the TME-associated cluster 4 displayed high expression of interferon-signature genes (e.g., IFIT1, IFIT2, IFIT3, MX1), suggesting that these may facilitate effector T cell function (Suppl. Fig. 3C).

To further define the genes and pathways dominant in the autologous MISTRG6 model, we utilized Ingenuity Pathway Analysis (IPA) to analyze pathways overrepresented in tumor compared with blood across cell types. We found significant associations with T cell activation in tumor including T Cell Receptor Signaling, Th1, Th2, Interferon Signaling, Jak/STAT Signaling, IL-17 Signaling, Antigen Presentation Pathways (Fig. 6A). In addition, genes representing the PD-1/PD-L1 Cancer Immunotherapy, T Cell Exhaustion Signaling, and Tumor Microenvironment Pathways were significantly associated with the TME.

**Figure 6:**
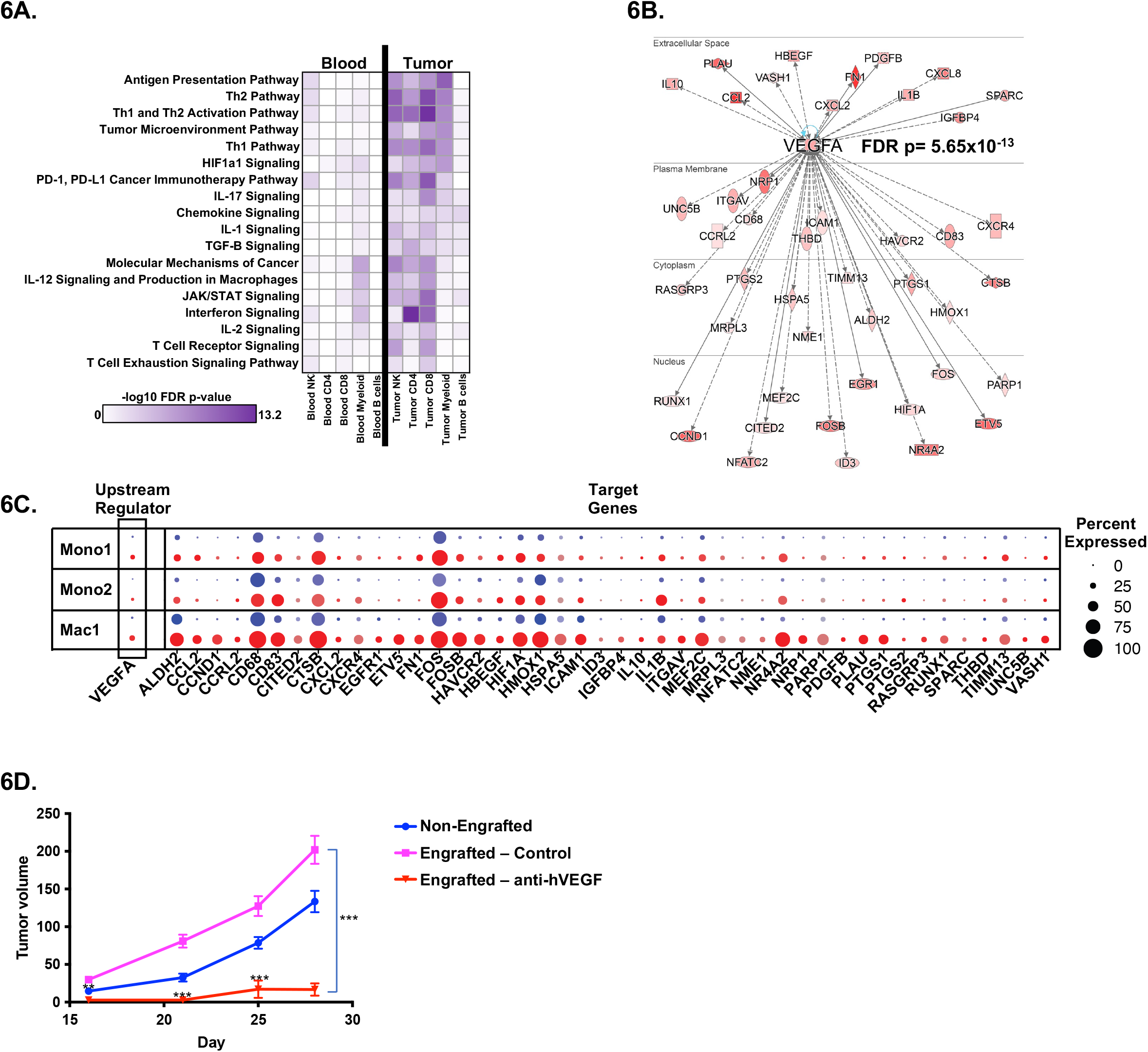
Immune cells in the TME display gene signatures associated with immune activation and signaling, including VEGF-A signaling, and blockade of this molecule abrogates enhanced tumor growth in engrafted autologous PDX mice. A. Canonical pathway representation in indicated cell types from blood (left) and tumor (right). B. Upstream pathway analysis identifies VEGFA and its target genes as highly represented in the TME; red shading indicates VEGFA target gene expression level in the TME. C. VEGFA and its target genes are over-represented in the TME (red) of myeloid gene clusters (monocyte 1, monocyte 2, macrophage 1); size of split dot plot circle indicates percent expression of each gene among cells of that cluster, while intensity of color indicates level of expression. D. Treatment of autologously engrafted MISTRG6 Mel2 PDX mice with bevacizumab, a clinical anti-hVEGF-A therapy, significantly reduces tumor growth compared with autologous HSPC-engrafted control (asterisks indicate comparison between drug-treated and control engrafted mice).

Notably, IPA Upstream Regulator Analysis identified VEGFA, a central player in tumor growth and vascularization across multiple tumor types,^18-20^ as a key upstream inducer of genes in the TME (FDR p= 5.65×10^−13^, Fig. 6B). Indeed, expression VEGFA itself was nearly absent in blood but induced in the TME, especially in macrophages (Fig. 6C). In addition, VEGFA direct and indirect targets were highly represented among the DEGs between tumor and blood in the myeloid cell types (Fig. 6B-C).

### Inhibiting the actions of human VEGF-A blocks the enhanced tumor growth in autologously engrafted mice

To test the imputed relevance of VEGF-A in the TME and its effect on tumor growth, we chose to selectively block human VEGF-A as the murine subcutaneous microenvironment is replete with murine VEGF which could compound the results. Thus, we treated autologous mice humanized with HSPCs and PDXs from Mel2 with the anti-human-VEGF-A antibody bevacizumab that has high affinity for human VEGF-A yet low affinity for mouse VEGF-A and which does not affect tumor growth in non-engrafted hosts.^10, 21^ The Mel2 PDXs grown in autologously engrafted MISTRG6 mice grew significantly larger than those in non-engrafted littermate control hosts. When treated with bevacizumab (10 mg/kg, weekly intraperitoneal injection), the enhanced tumor growth seen with human autologous engraftment was significantly abrogated, with bevacizumab-treated mice bearing significantly smaller tumors compared with controls (Fig 6D). Thus, these *in silico* and *in vivo* results suggest that human VEGF-A production in the TME of mice bearing an autologous human TME enhances tumor growth in MISTRG6 PDX models.

## Discussion

While advances in immunotherapy have greatly improved quality and length of life for people with cancer, there remains significant variability in individual patient responses to treatment. For example, nearly 40% of patients with metastatic melanoma, a disease with among the most immunotherapy successes, do not respond initially to combination CTLA-4 and PD-1 inhibition and of those that respond, the duration of response is limited by as-yet undefined mechanisms of acquired resistance.^22, 23^ Hence, there is a need for personalized models that faithfully reflect the TME of individual patients, and that can therefore be used to test primary and acquired resistance to therapies. By engrafting mice with bone marrow derived stem cells followed by implantation of tumor derived from the same donor, we have demonstrated that autologous MISTRG6 models recapitulate important features of the human TME, including sufficient immunosuppression to prevent tumor clearance, presence of activated/exhausted-like T cells, and harboring innate immune cells including DCs, monocytes, NK cells, and macrophages, the latter being especially relevant by the production of VEGF-A.

While humanized mice have long been utilized for engraftment of human hematopoietic cells to study myriad processes,^24^ MISTRG6 mice allow engraftment with few HSPCs and creation of humanized mice that reflect the immune makeup of living patients. Indeed, we found that a low-volume BM aspiration can provide sufficient HSPCs (1-5 × 10^6^) to engraft multiple mice on a scale not seen previously. These cohorts are large enough to facilitate drug or other comparisons, opening the door to exciting experiments such as prospective co-clinical trials.

Importantly, MISTRG6 mice engrafted autologously with BM-derived patient HSPCs support tumor growth for multiple weeks, permitting study of tumor expansion in a timeline similar to other models. Notably, we found that tumors grown in autologously-engrafted hosts were larger than those grown in non-HSPC-engrafted hosts, underscoring the functionality of the human cell engraftment and the pro-growth effects that the TME can acquire and provide. However, not all autologous models displayed enhanced growth in the presence of autologous human cells, underlining the inter-patient differences that our MISTRG6 model elucidates and that will be important for future personalized medicine approaches.

The ability to study the myeloid cell fraction of the TME is critical to understanding tumor-immune interactions. Indeed, myeloid lineage cells, such as macrophages and dendritic cells, are major components of a patient’s TME that are recapitulated in our approach, and these innate immune cells have drastic and varied effects on tumor growth as well as immune function^25, 26^ Our single cell transcriptomic analyses pointed to the myeloid compartment as producers of VEGF-A in the TME, a central pro-tumor cytokine. In support of this cytokine’s importance, functional inhibition of human VEGF-A with bevacizumab abrogated the enhanced growth effects in the model and indicates the potential of the model for pre-clinical drug testing.

Myeloid cells are also important orchestrators of adaptive immune responses, functioning as antigen-capturing and -presenting cells, as well as modulators of the activity of other immune cells, especially effector T cells. Each of these cell types can have direct pro- and anti-tumor effects on malignant cells. For example, macrophages can exhibit pro- or anti-inflammatory phenotypes. The former are linked with type 1 inflammation, intracellular pathogen killing, and tumor resistance, while the latter exhibit immunoregulatory and pro-angiogenic properties, promote tissue repair, and contribute to tumor progression^3, 27-30^. In this regard, the demonstration of several macrophage subtypes in the TME of autologously engrafted MISTRG6 mice may provide a way to decipher these processes *in vivo*. Finally, the presence of DCs in our model makes it more likely that the T cell effector functions that occur have biologic relevance since they can occur in an MHC-matched manner.

Thus, due to their unique sensing abilities, their functional plasticity, and their central position in the TME, macrophages, monocytes, and DCs are essential elements to investigate the variability in response to cancer and to its treatment. It is critically important to credential a model of humanized mice where these lineages are properly developed and functionally mature.

Like other pre-clinical models, our method has limitations. While MISTRG6 autologous mice develop major innate immune cell types as discussed, like most other humanized mouse models, they lack large numbers of human neutrophils, basophils, and eosinophils; further research is required to identify the genes required to support the development of these human cell types. In addition, like other PDX methods, the autologous technique is limited by the fact that not all fresh tumor tissue samples generate efficient PDXs that grow robustly in the timeline of an experiment. In addition, the yield of BM aspiration is variable, and in rare cases too few HSPCs are obtained. Further humanization of recipient mice is likely to improve engraftability, such that autologous mice may be engrafted with fewer and fewer adult HSCs. Finally, the chronological lag between tissue collection and autologous modeling inherent in the technique due to the need for PDX generation, which can take weeks or months, may limit the ability to use the model as a predictor of primary therapies. However, given that surgical tissue is often collected early in a patient’s clinical course (e.g., for resection of a primary NSCLC), autologous modeling can begin early in a patient’s course, and bear fruit in modeling second or third therapies. Furthermore, PDX models can be successfully generated from small biopsy specimens, broadening the applicability to most metastatic solid tumors^31-33^. Our findings demonstrate the broad utility of a genetically matched, fully autologous humanized mouse model system for investigating the TME and treatment responses of individual solid tumor patients.

## Acknowledgements

This work was funded by the Howard Hughes Medical Institute (R.A.F.). This study was also supported, in part, by National Institute of Health grants T32HL007974 (M.C.), K08CA245211 (M.C.), R01CA248277 (R.C.F and R.A.F.), T32CA009621 (K.J.R and B.K.). M.C. and R.A.F. received funding from the Yale SPORE in Lung Cancer 1P50CA196530-01. R.C.F received funding from The Alvin J. Siteman Cancer Center Siteman Investment Program, The Foundation for Barnes-Jewish Hospital Cancer Frontier Fund, the National Cancer Institute Cancer Center Support Grant P30 CA091842, and the Barnard Trust. R.C.F. also received funding from the Washington University PDX Development and Trial Center (U54CA224083) and the David Riebel Cancer Research Fund. We thank the Siteman Tissue Procurement Core and the core grant/services of the Washington University Digestive Diseases Research Core Center (P30 DK052574) for supporting this work. We are grateful to Ricky Brewer, Eleanna Kaffe and other members of the Flavell laboratory for helpful discussions and comments; J. Alderman, C. Lieber, B. Cadugan, C. Hughes, D. Urbanos and E. Hughes-Picard for administrative and technical assistance; P. Ranney, C. Weibel and for mouse colony management; H. Lazowski, C. Bensley and A. Wurtz for assistance with patient consent; L. Devine for assistance with cell sorting; G. Wang of the Yale Center for Genome Analysis for assistance with scRNAseq experiments. This article is subject to HHMI’s Open Access to Publications policy. HHMI lab heads have previously granted a nonexclusive CC BY 4.0 license to the public and a sublicensable license to HHMI in their research articles. Pursuant to those licenses, the author-accepted manuscript of this article can be made freely available under a CC BY 4.0 license immediately upon publication.

## Contributions

M.C. conceived the project, collected patient samples, performed experiments, interpreted data and wrote the manuscript. J.M. and B.K. performed experiments, interpreted data and edited the manuscript. Y.Z., K.J.R, R.Q., G.K., Z.S., L.H., F.B., A.A., J.Z., L.S., E.S., J.M., Y.B., A.O. performed experiments and interpreted data. S.P.G., M.G., O.G., S.F., R.G.M, Y.K. interpreted data. D.B., F.D., A.D., J.B., B.J., S.G., K.P. assisted with patient consenting and tissue collection. A.K.P. and R.C.F conceived the project, interpreted data and edited the manuscript; R.A.F. conceived and oversaw the project, interpreted data and edited the manuscript.

## Supplementary Figure Legends

**Supplementary Figure 1:**
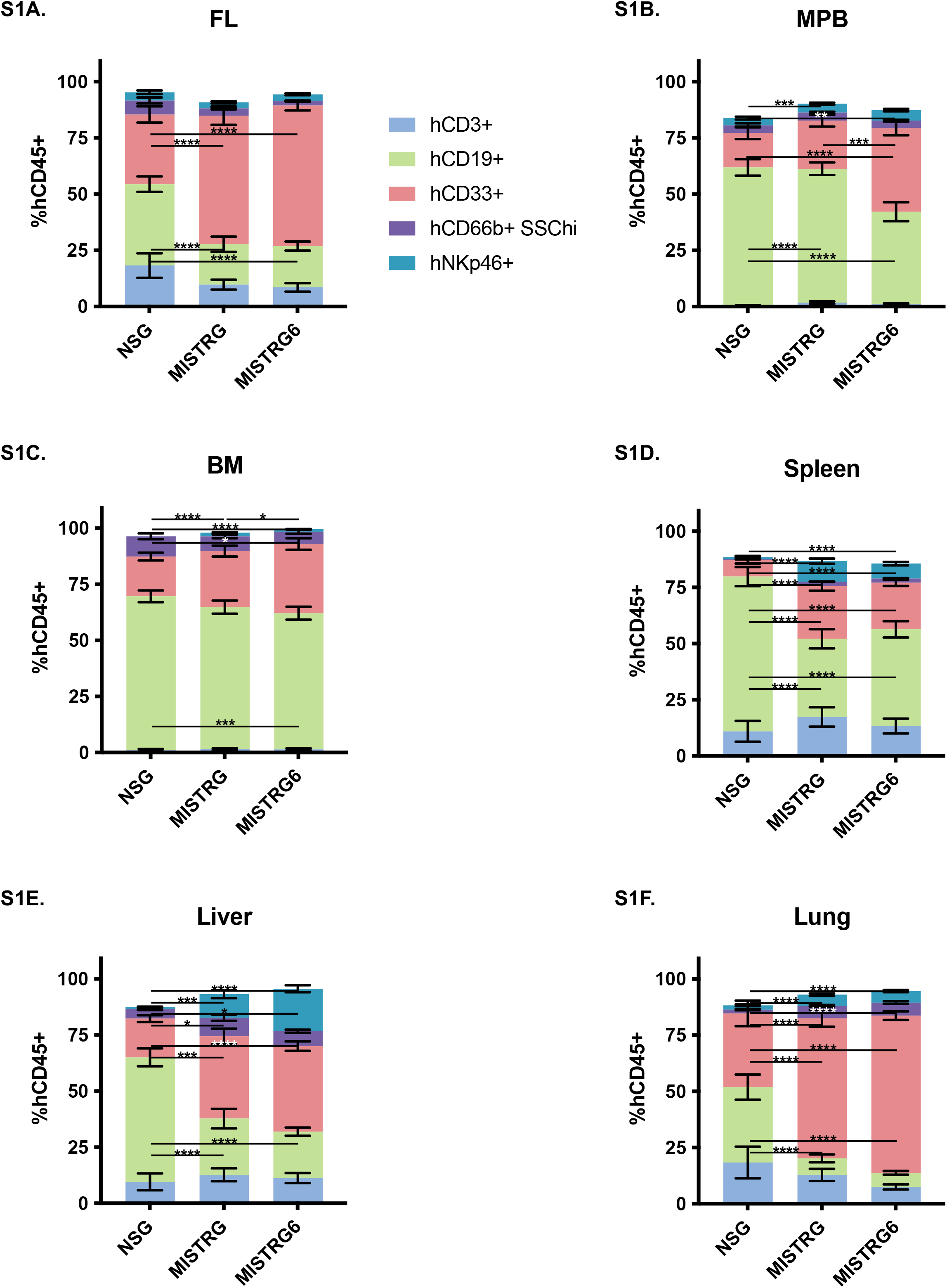
MISTRG6 mice display robust peripheral blood and tissue engraftment of human cell types, including innate immune cells. A. Human hematopoietic cells in peripheral blood from mice of indicated strains engrafted with FL-derived CD34^+^ cells, including T cells (hCD3^+^, blue), B cells (hCD19^+^, green), myeloid cells (hCD33^+^, red), neutrophils (hCD66b, SSC^hi^, purple), NK cells (hNKp46^+^, teal). B. Human hematopoietic cell profiles in peripheral blood from mice of indicated strains engrafted with CD34^+^ cells isolated from mobilized peripheral blood (MPB) of adult donors. C. Human hematopoietic cell profiles in BM from mice of indicated strains engrafted with FL-derived CD34^+^ cells. D. Human hematopoietic cell profiles in spleen from mice of indicated strains engrafted with FL-derived CD34^+^ cells. E. Human hematopoietic cell profiles in liver from mice of indicated strains engrafted with FL-derived CD34^+^ cells. F. Human hematopoietic cell profiles in lung from mice of indicated strains engrafted with FL-derived CD34^+^ cells.

**Supplementary Figure 2:**
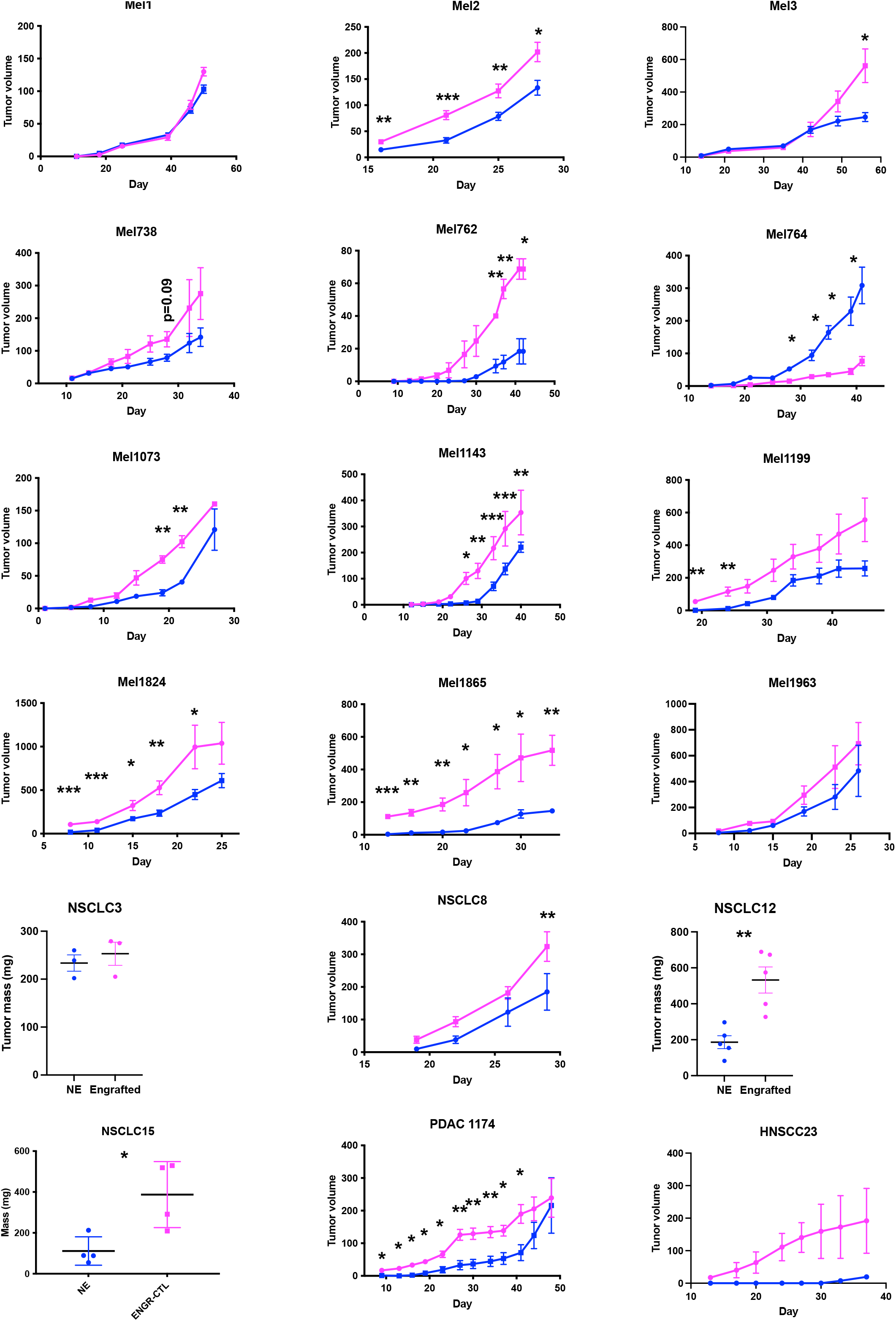
Autologously-engrafted MISTRG6 mice support enhanced tumor growth in multiple patient models. Tumor growth curves for non-engrafted (blue) and autologously engrafted (magenta) PDX mice representing indicated patients; * p<0.05, ** p<0.01, *** p<0.001, unpaired parametric t-test; bars indicate mean and S.E.M.

**Supplementary Figure 3:**
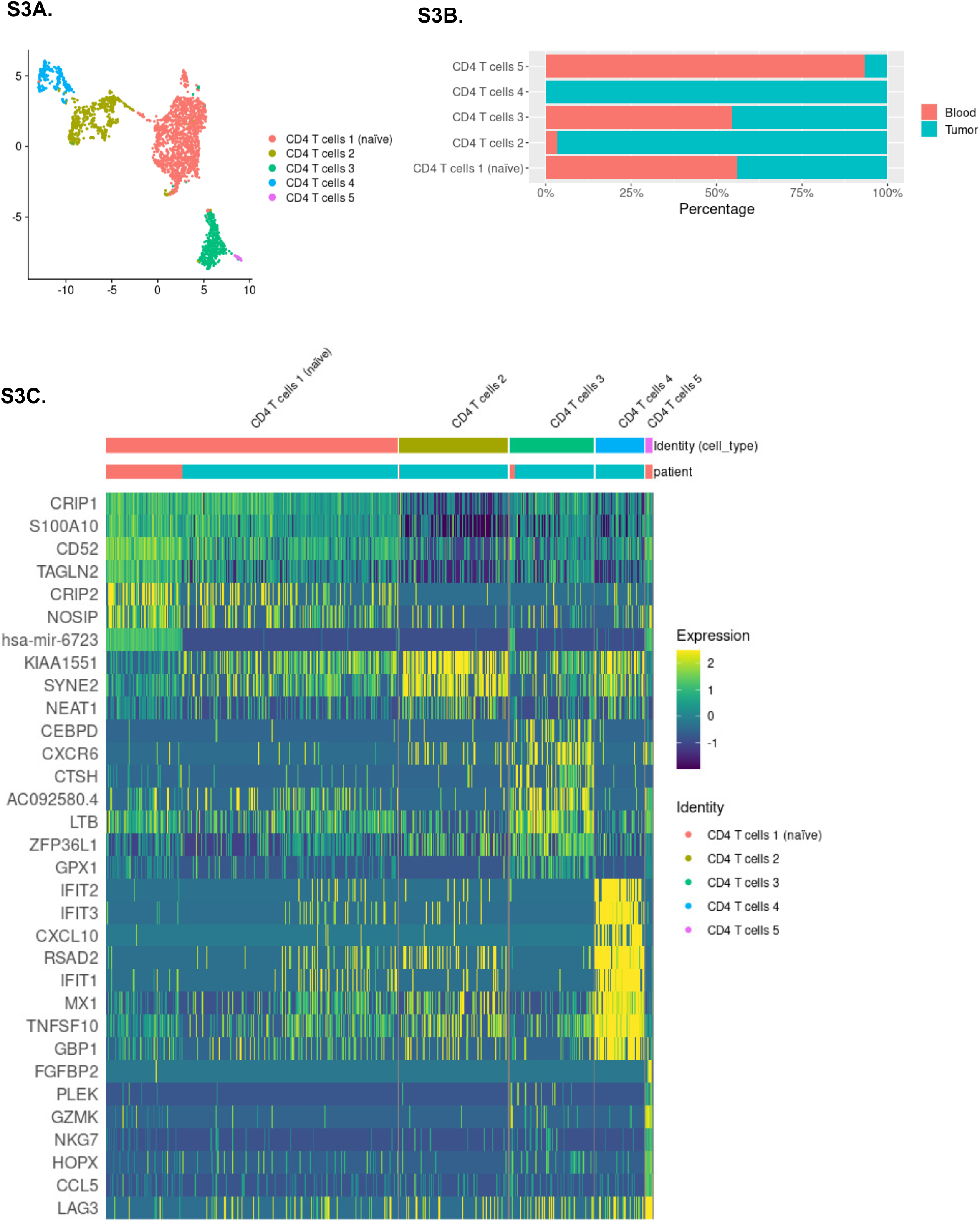
Blood and tumor-resident CD4 T cells display distinct transcriptional states including naïve and interferon-responsive subtypes. A. Re-clustering of CD4 T cells reveals sub-structure of 5 clusters. B. Cluster representation of CD4 T cell subclusters in tissues. C. Heatmap indicating expression of top differentially expressed genes between each cluster, highlighting interferon activation signature of CD4 4 cluster.

**Supplementary Table 1.**
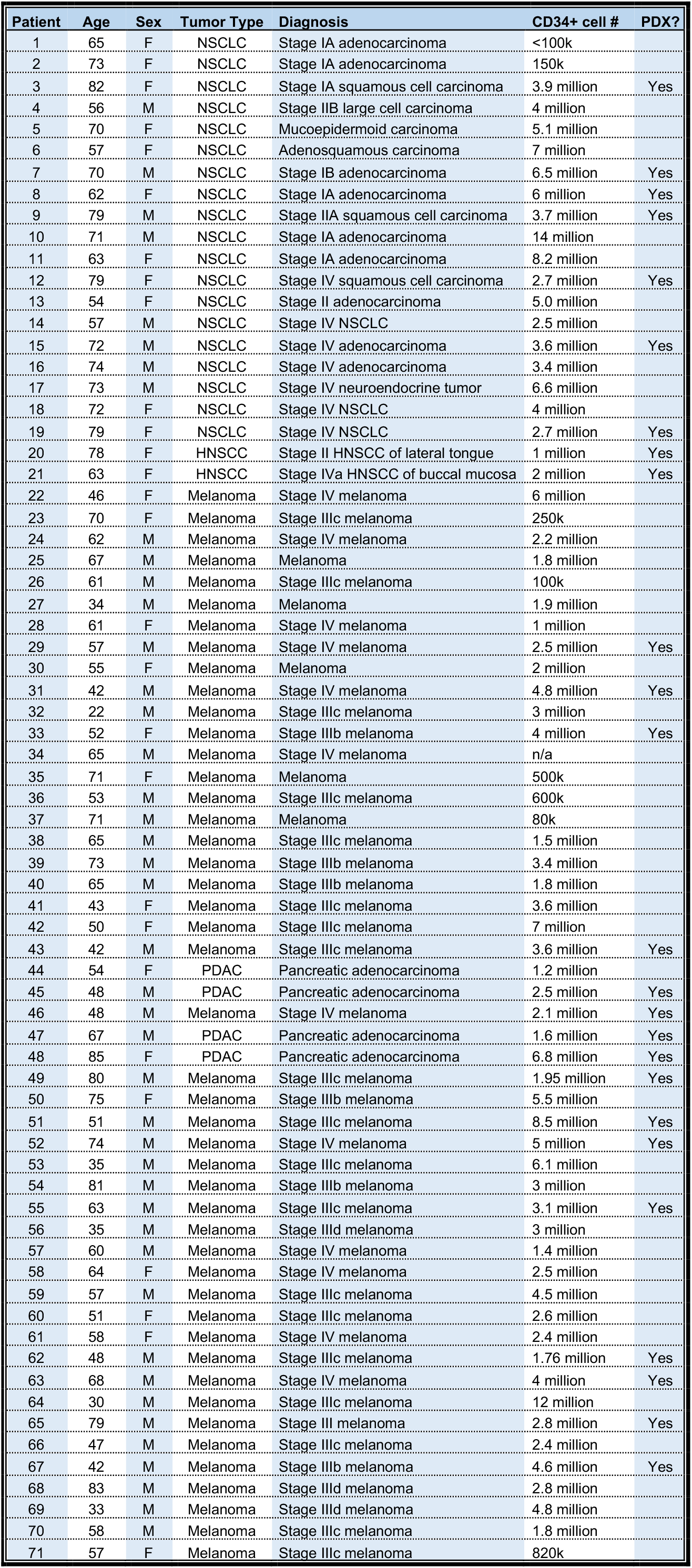
Clinical data of patients enrolled in prospective study collecting tumor and BM from solid tumor patients; indicated PDXs grew sufficiently for autologous humanized mouse modeling.

## Methods

### Mice

MISTRG6 mice bear human knock-ins of M-CSF (CSF1), CSF2/IL3, SIRPA, THPO genes were humanized on a RAG2^-/-^, IL2RG^-/-^ immunodeficient background, as previously described ^10, 13, 14^. Mice were maintained in a BSL-2 mouse facility on a 14-10 hr light-dark cycle with 2 weeks on, 2 weeks off sulfamethoxazole/trimethoprim diet. Experimental mice were cross-fostered with CD1 female mice from Charles River Laboratories for maintenance of eubiotic flora. Newborn mice engrafted with adult HSPCs were pre-conditioned with 150 rads of irradiation using an X-ray irradiator prior to intrahepatic injection as previously described^10^.

### Human HSPC Isolation and Enrichment

All human samples were obtained and handled in accordance with established Yale and Washington University Human Investigational Committee protocols. Candidates for participation were identified by investigators, and subsequently approached by investigators or clinical trials staff to discuss participation in this study. Patients were prospectively enrolled in Yale HIC #1603017380 or WUSM IRB # 201108117 after informed consent. FL HSPCs were purified as previously described^34^. BM aspirates were performed using a 15 Gauge Illinois needle (VWR) with standard technique. 40-150 cc of BM was aspirated from unilateral or bilateral iliac crests. Peripheral blood was also collected from an intravenous line placed in the normal course of perioperative care. The BM aspirate, peripheral blood, and a component of the resected tumor were subsequently transported to our laboratory for further processing. Freshly-aspirated BM and peripheral blood were anticoagulated with sterile EDTA and subjected to Ficoll density-mediated separation for isolation of mononuclear cells. CD34+ cells were positively selected using the EasySep Human CD34 Positive Selection Kit (STEMCELL Technologies) according to the manufacturer’s protocol. HSPCs and PBMCs were cryopreserved in 90% FBS + 10% DMSO until use.

### Tissue Processing and Flow Cytometry

Fresh excess human tumor tissue was obtained from enrolled patients and divided into 1-2 mm^3^ pieces for subcutaneous implantation in the flank of MISTRG6 or NSG mice lacking human HSPC reconstitution, as previously described^31, 35^. Tumor growth was monitored by caliper measurements, and when tumors reached 500-1000 mm^3^, mice were euthanized and tumor tissue again divided into 1-2 mm^3^ pieces for PDX passaging. PDXs were passaged 2-3 times, and histology confirmed with H&E staining. PDXs were cryopreserved in 90% FBS + 10% DMSO in liquid nitrogen until use.

Tumors, spleen, lung, and liver tissues were minced in PBS+1%FBS and digested with collagenase D (1mg/ml, Sigma) and DNAse at 37°C for 15 minutes. Tumor-infiltrating immune cells were isolated with a 40-80% Percoll density gradient (Sigma). Non-specific staining was blocked with human (BD Biosciences) and mouse (Bio X Cell, BE0307) Fc block for 10 min. Epitopes were stained at 4°C for 30 minutes and cells were fixed with 4% paraformaldehyde; intracellular staining was performed using BD permeabilization buffer (BD Biosciences). Samples were analyzed on an LSRII flow cytometer, and data parsed with FlowJo software v10 as well as GraphPad Prism v9. Cell sorting was performed using a BD FACS Aria flow sorter under BSL2 containment. Sorted cells were subjected to 10X Genomics 5’V2 library construction and sequenced using NovaSeq. Data were filtered as described previously^34^.

### Immunofluorescence staining

Cryosections (8 um) were acetone fixed, air dried, washed with PBS and consecutively treated with Fc Receptor Block (Innovex Bioscience) for 40 min + Background Buster (Innovex Bioscience) for an additional 30 min. The sections were then stained with directly conjugated antibody mix in PBS 5%BSA 0.1%Saponin for 1 hour at room temperature and washed. Nuclei were counterstained with SytoxBlue 1:1000 for 2 min. Tissues were mounted in Fluoromount-G mounting media. Images were acquired using a Leica SP8 confocal microscope.

### Statistical Analyses

Unpaired parametric t-tests were utilized to compare immune cell frequencies and tumor sizes, except where indicated otherwise in figure legends. Bars indicate mean +/- S.E.M., except where indicated otherwise. The single-cell RNA-seq data analysis was performed using Seurat v4.0.1 R package,^36^ including cell type stratification and comparative analyses between tumor and blood samples. In the quality control (QC) analysis, poor-quality cells with < 250 (likely cell fragments) or > 5,000 (potentially doublets) unique expressed genes were excluded. Cells were removed if their mitochondrial gene percentages were over 25% or if their ratios of reads mapped to human genome (over the total reads mapped to humanized mouse genome) were lower than 90%. The data was first integrated with default settings in Seurat (using 30 dimensions in the anchor weighting procedure), followed by principal component analysis (PCA) for dimensionality reduction. We retained 30 leading principal components for further visualization and cell clustering. The Uniform Manifold Approximation and Projection (UMAP) algorithm was used to visualize cells on a two-dimensional space.^37^ Subsequently, the share nearest neighbor (SNN) graph was constructed by calculating the Jaccard index between each cell and its 20-nearest neighbors, which was then used for cell clustering based on Louvain algorithm (with a resolution of 0.5). Each cluster was screened for marker genes by differential expression analysis based on the non-parametric Wilcoxon rank sum test. Based on checking the expression profile of those cluster-specific markers, we identified 16 distinct cell types, including different subtypes of CD4/CD8 T cells, B cells, myeloid cells, NK cells, and melanoma cells. In the downstream analysis, we focused on myeloid cells, CD8 T cells, and CD4 T cells respectively. Specifically, we separated out each cell type and redid the clustering to define cell subtypes (states) in a high-resolution manner. Differential expression analysis was performed to 1) reveal the functional role of each subtype, and 2) identify differentially expressed genes (DEGs) between the blood and tumor conditions. Top representative DEGs were visualized using heatmaps or dot plots. For Ingenuity Pathway Analysis, differentially expressed genes were identified by filtering group comparisons for biological (ABS(FC) > 1.5) and statistical significance (adjusted p < 0.05). These were uploaded into Ingenuity Pathway Analysis (IPA) software (Qiagen, Content version 73620684, 2022 http://www.ingenuity.com), mapped to their corresponding IPA identifier, and used for functional analysis. Pathways and upstream regulators were determined using an FDR p <0.05 calculated with a right-tailed Fisher’s exact test.

### Sequence Alignment

WES data were processed using the immuno.cwl analysis workflow (https://github.com/genome/analysis-workflows/tree/master/definitions/pipelines). Briefly all alignment data was processed through xenome (1.0.0) and reads classified as either “graft” (*H. sapiens*) or “both” were retained in order to remove incidental data from the host xenograft samples. WES data was then aligned to GRCH38 via bwa-mem (0.7.15) and read duplicates marked with picard (2.18.1). BQSR correction was performed via GATK (4.1.8.1). Variant calling was performed via Mutect (GATK4.2.3.0), Strelka (2.9.9), Varscan2 (2.4.2), and Pindel (1.4.2), and a log-likelihood ratio filter applied. Called variants were further refined by the following criteria: variant caller count > 2, tumor depth >= 50 & tumor variant count >=2, normal depth >= 20 & normal variant count <=3. Variant intersection amongst related samples was then performed using R (4.1.1) and Upset plots created using the UpSetR R package.

### Antibodies

mCD45 (Clone: 30-F11), hCD45 (HI30), hCD3 (UCHT1), hCD14 (HCD14), hCD16 (3G8), hCD19 (HIB19), hCD33 (WM53), hCD20 (2H7), hCD11B (M1/70), hCD11C (3.9), HLA-DR (LN3), hPD1 (A17188B), hNKp46 (9E2), hCD4 (OKT4), hCD8 (SK1), mCD117 (2B8), hCD117 (104D2), mSca1 (D7), hCD34 (581), hCD38 (HIT2), mCD34 (HM34), mCD38 (90), hCD31 (WM59). All antibodies were obtained from BioLegend and used at 1:300 dilution in PBS + 1% FCS.

